# Behavioral and neural signatures of working memory in childhood

**DOI:** 10.1101/659409

**Authors:** Monica D. Rosenberg, Steven A. Martinez, Kristina M. Rapuano, May I. Conley, Alexandra O. Cohen, M. Daniela Cornejo, Donald J. Hagler, Kevin M. Anderson, Tor D. Wager, Eric Feczko, Eric Earl, Damien A. Fair, Deanna M. Barch, Richard Watts, BJ Casey

## Abstract

Working memory function changes across development and varies across individuals. The patterns of behavior and brain function that track individual differences in working memory during development, however, are not well understood. Here we establish associations between working memory, cognitive abilities, and functional MRI activation in data from over 4,000 9–10-year-olds enrolled in the Adolescent Brain Cognitive Development study, an ongoing longitudinal study in the United States. Behavioral analyses reveal robust relationships between working memory, short-term memory, language skills, and fluid intelligence. Analyses relating out-of-scanner working memory performance to memory-related fMRI activation in an emotional *n*-back task demonstrate that frontoparietal activity in response to an explicit memory challenge indexes working memory ability. Furthermore, this relationship is domain-specific, such that fMRI activation related to emotion processing during the emotional *n*-back task, inhibitory control during a stop-signal task, and reward processing during a monetary incentive delay task does not track memory abilities. Together these results inform our understanding of the emergence of individual differences in working memory and lay the groundwork for characterizing the ways in which they change across adolescence.

## Introduction

Working memory—a collection of cognitive processes responsible for storing and manipulating information—is a foundational ability that varies widely across individuals. Individual differences in working memory, which appear to be stable over time (1–5), have pronounced real-world significance. Although the direction of causality is unclear, working memory explains approximately 20–30% of the variance in fluid intelligence in children (6) and more than 40% of this variance in adults (7). Furthermore, working memory function, which is related to executive and visuospatial attention (8, 9), short-term memory (10), and inhibitory control (11), predicts consequential outcomes in development, including reading and math skills (12–15). Despite the theoretical and practical importance of characterizing associations between working memory and other mental processes, much remains to be learned about the nature of these relationships in the developing and developed mind.

Working memory not only varies across individuals, but also changes across the lifespan. Working memory emerges in infancy and develops rapidly over the first year of life (16–19). It continues to improve during childhood, plateaus in mid-to-late adolescence (20–24), and declines after age 40–50, albeit less steeply than it changed in early development (25–28). Developmental gains in working memory follow improvements in attention shifting, attentional maintenance, and distractor suppression (18), whereas changes during later childhood accompany increases in domain-general processing speed and memory capacity (29–31) with developmental asymptotic performance by adolescence (32–35). Decrements in older adulthood relate to declines in processing speed, selective attention, and distractor suppression (36–38).

Importantly, converging neuroimaging evidence suggests that variation in frontoparietal brain systems, involved in processes including attention and cognitive control (39–41), accounts for both developmental change in working memory and individual differences in working memory in adulthood. Early work demonstrated that the same middle and inferior frontal regions that support working memory performance in adults also support performance in children (42). This evidence led to theorizing that the protracted fine-tuning of prefrontal circuitry contributes to working memory improvements during childhood and adolescence (33, 34). Longitudinal studies support this prediction, with evidence that maturation in prefrontal and parietal volume and structural connectivity accompany working memory development (43, 44). Cross-sectional work suggests that increases in frontoparietal activation during working memory tasks are associated with age-related improvements in performance (32, 45–47). In the developed brain, individual differences in frontoparietal areas’ microstructure, function, and structural and functional connectivity track individual differences in working memory (48–52). A subset of developmental studies show similar associations between in-scanner working memory performance (a state-like measure of memory function) and frontoparietal activity during working memory tasks when controlling for age (45, 47). However, it is not yet known whether frontoparietal network function during memory challenges—or during cognitive task challenges more generally— predicts individual differences in working memory during development.

Here we examine the emergence of behavioral and neural signatures of working memory in childhood. Using data from more than 4,000 9–10-year-olds participating in the Adolescent Brain Cognitive Development (ABCD) study (53, 54), we first establish relationships between working memory and other cognitive and attentional abilities, including short-term memory, language and verbal skills, fluid intelligence, processing speed, attention, inhibitory control, and reward processing. Because the ABCD study will follow children longitudinally for ten years, characterizing these associations in childhood not only informs the structure of cognition at a single time point, but also facilitates understanding the ways in which this cognitive structure changes across adolescence. We next ask whether performance on an out-of-scanner working memory test is related to frontoparietal brain activity when measured *(a)* during a working memory challenge and *(b)* during task challenges unrelated to memory. Together our results provide insight into the emergence of individual differences in working memory, and underscore the importance of task fMRI as a “stress test” for cognition (55) that can reveal task-specific and task-general neural signatures of a mental process or behavior.

## Results

### Working memory performance in childhood

Individual differences in working memory and other cognitive and attentional processes were assessed using data from 4398 9–10-year-olds in the Adolescent Brain Cognitive Development (ABCD) study, an ongoing 21-site longitudinal study of neurocognitive development (56). ABCD study data collection includes yearly behavioral assessments, interviews, questionnaires, and biosample collection as well as biennial MRI scans (54). Year-one (baseline) demographic, behavioral, and functional MRI data from the first half of the cohort are analyzed here.

Working memory performance in the ABCD cohort, measured with the National Institutes of Health (NIH) Toolbox List Sorting Working Memory Test, approximated the normative population mean (uncorrected standard score mean = 98.2, *s.d.* = 11.2, range = 40–136; normative mean = 100, *s.d.* = 15). Working memory was positively correlated with age (*rs* = .13, *p* < 2.2×10^−16^) and numerically differed by sex, albeit with a negligible effect size (female mean = 97.8; male mean = 98.5; Welch *t*_4339.8_ = 1.94; *p* = .052; Cohen’s *d* = .06). Performance on all other neurocognitive measures in the ABCD task battery—which assess short term memory, fluid intelligence, visuospatial attention, reading and language skills, cognitive control, processing speed, flexible thinking, learning, delay of gratification, emotion regulation, impulsivity, and reward processing—is visualized in Fig. 1.

**Figure 1.**
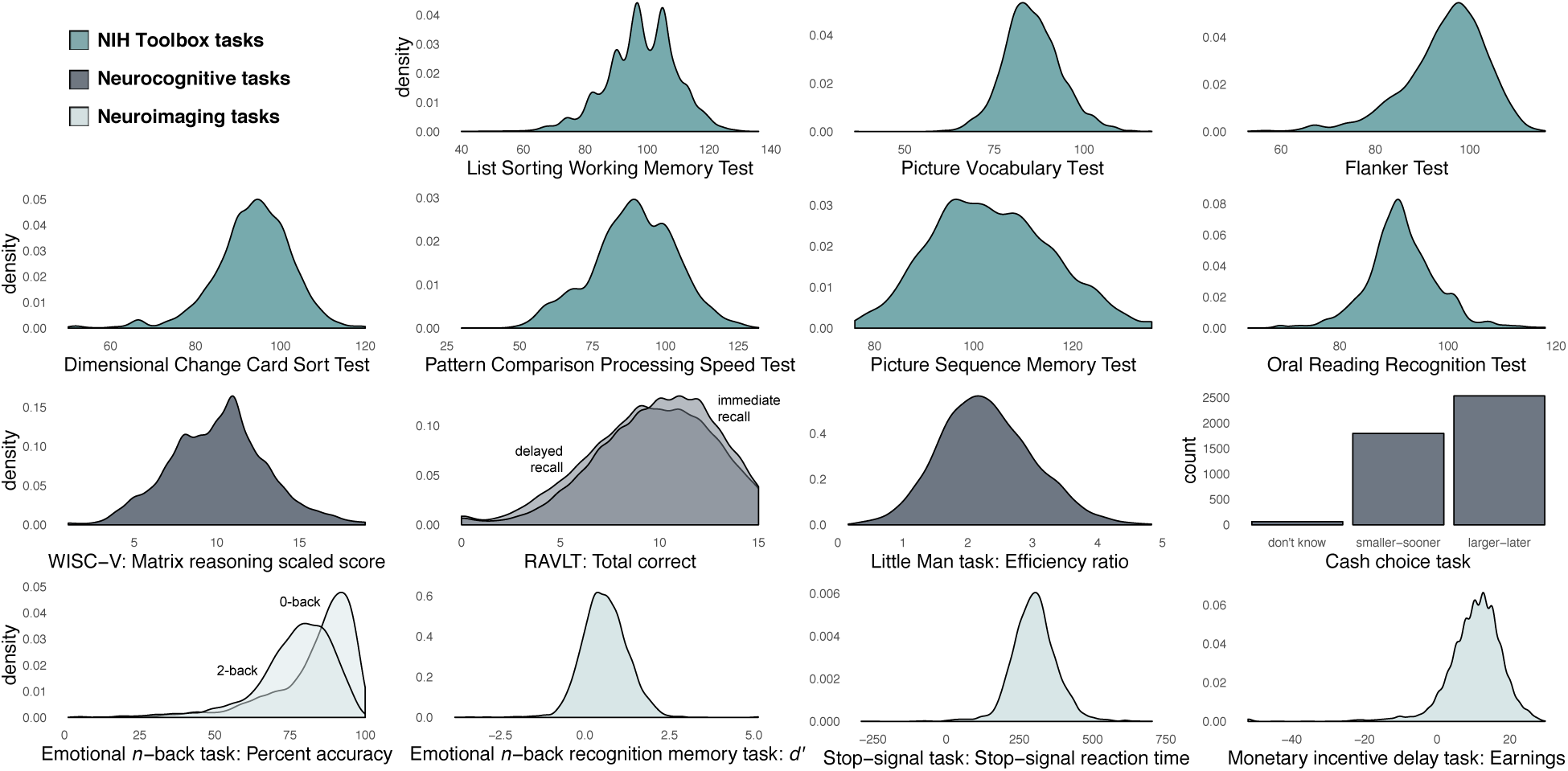
Kernel density estimates, or smoothed histograms, show performance in the full sample of 4,398 9–10-year-olds, including statistical outliers. NIH Toolbox performance is measured with uncorrected standard scores. Responses on the cash choice task—whether a child preferred to receive a smaller–sooner reward, a larger– later reward, or couldn’t choose—are visualized with a histogram. Although “don’t know” responses on this task are included here, they were excluded from formal analysis.

### Behavioral signatures of working memory

Although a rich literature in cognitive psychology describes relationships between working memory and cognitive and attentional processes in adulthood, how these associations emerge in development is less well understood. Thus, a primary goal of the current work is to characterize these associations in childhood in order to understand how they change across development.

To relate working memory to cognitive and attentional abilities, we computed pairwise Spearman correlations between performance scores on all tasks included in the dataset (i.e., all behavioral measures visualized in Fig. 1). Correlation coefficients are reported without corresponding *p*-values because effect sizes as small as *r*^2^ = .0009 are significant at *p* < .05 in a sample of 4398, and statistical dependence introduced by family relatedness, site effects, and the inclusion of multiple performance measures per test precludes parametric *r*-to-*p* conversion. Furthermore, the goal of this analysis is to establish a pattern of behavioral relationships rather than to evaluate the statistical significance of particular associations.

Across individuals, working memory was most strongly related to language skills measured with the NIH Toolbox Oral Reading Recognition and Picture Vocabulary tests (*rs* values = .37); memory-related performance on the emotional *n*-back task and Rey Auditory Verbal Learning Test (RAVLT); and fluid intelligence measured with matrix reasoning (*rs* = .32; Figs. 2, 3).

**Figure 2.**
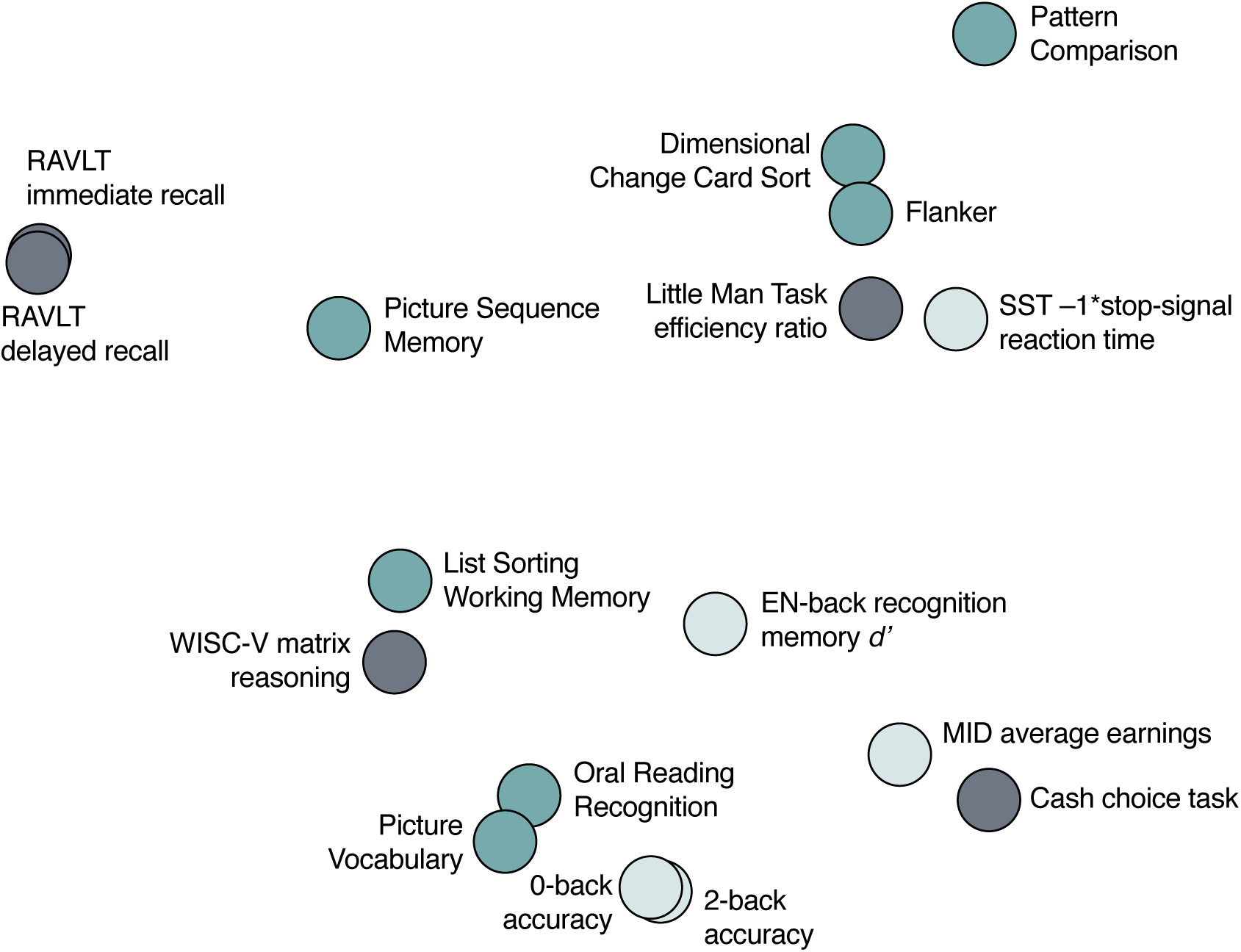
Multidimensional scaling plot illustrating two-dimensional distance between behavioral metrics in children with no missing data (*n* = 2,304). Classical multidimensional scaling was applied to the complete-case sample to avoid assumptions associated with imputing missing values. Distance was calculated as the Euclidean distance between each pair of behavioral measures after mean-centering and scaling each measure across participants. NIH Toolbox measures are shown in dark green, other neurocognitive measures in dark gray, and neuroimaging task measures in light green.

**Figure 3.**
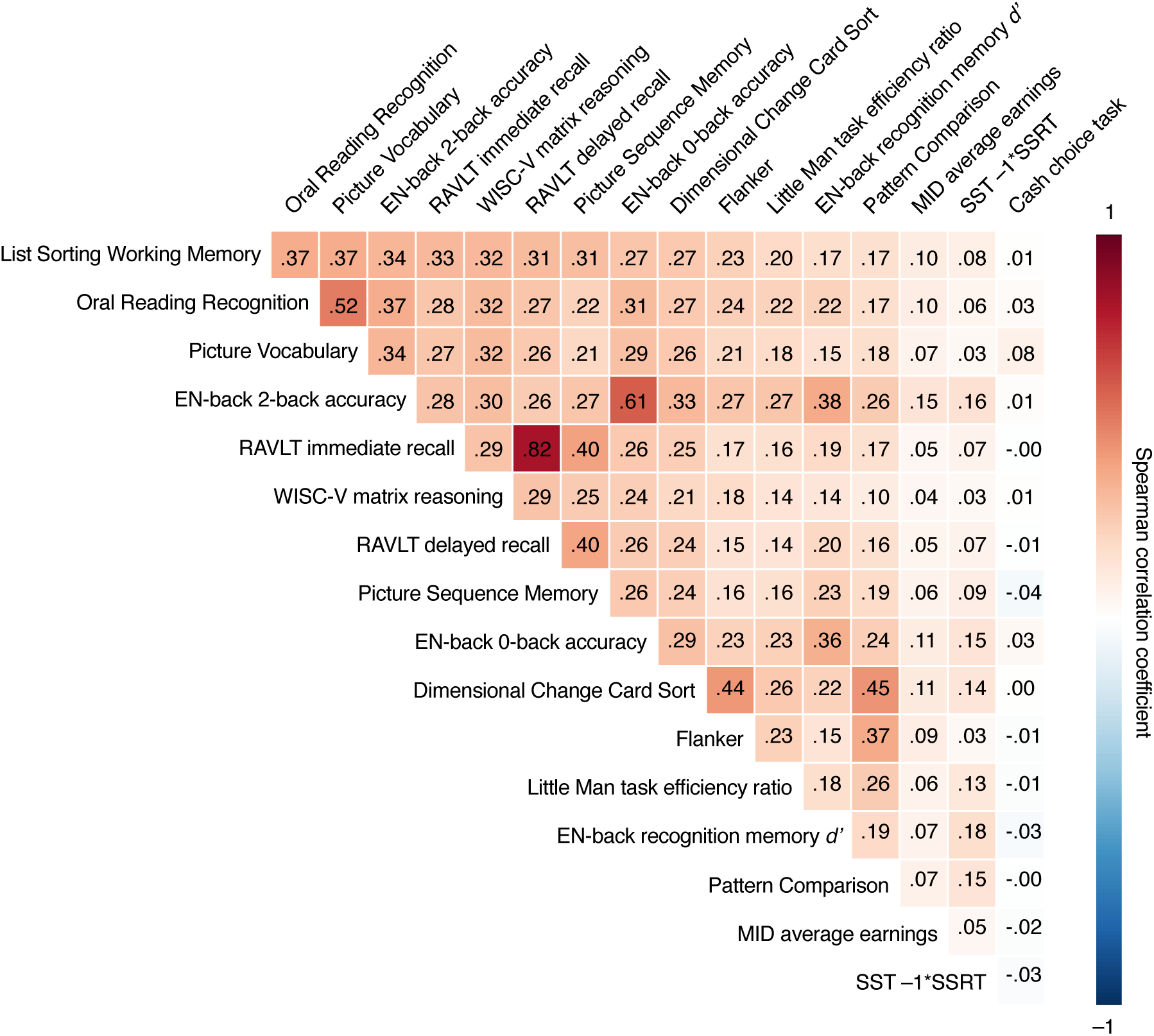
Spearman correlations between performance measures in the full 4,398-child sample. Measures are ordered according to the strength of their relationship with working memory, operationalized as NIH Toolbox List Sorting Working Memory Test. Because the outcome of the cash choice task is binary, relationships with performance on this measure are equivalent to point-biserial Spearman correlation coefficients.

Correlations between list sorting performance and performance on other memory tests also revealed relationships between different aspects of memory. The emotional *n*-back task, collected during functional MRI, measured performance during high memory load (2-back) and low memory load (0-back) task blocks. During 2-back blocks, children were asked to indicate when they saw a picture identical to the one they saw two trials back. During 0-back blocks, children were shown a target picture and instructed to indicate when they saw a matching image. Working memory was more strongly related to 2-back than to 0-back accuracy (*rs* = .34 vs. .27; Steiger’s *z* = 5.10, *p* < .001), indicating that, as predicted, working memory ability is reflected to a greater degree by performance on high-load vs. low-load *n*-back blocks. Recognition memory for emotional *n*-back stimuli (happy, fearful, and neutral face photographs and place photographs) was tested after fMRI data collection. Although working memory was less highly correlated with recognition memory than with performance on visual attention tasks, including the Flanker (*rs* = .17 vs. .23; Steiger’s *z* = 3.27, *p* = .001) and Little Man (*rs* = .17 vs. .20; Steiger’s *z* = 1.45, *p* = .15) tasks, this may reflect low overall memory for specific stimuli, especially face photographs, at this age (54). Finally, the RAVLT assessed immediate recall of a word list and list recall after a 30-minute delay. Working memory was numerically more closely related to immediate than to delayed recall on this task (*rs* values = .33 vs. .31; Steiger’s *z* = 1.62, *p* = .10), distinguishing relationships between working memory and short-term memory across shorter and longer delays.

### Behavioral relationships are not influenced by family structure

The full 4398-child cohort includes 1149 related children from 3819 unique families (based on self report). Because relatedness affects the independence of behavioral measures and thus could have affected relationships between them, we replicated correlations between cognitive and attentional abilities in a subset of data from only one child per family (Supplementary Fig. 1). The pattern of behavioral relationships in this unrelated subsample was nearly identical to that observed in the full sample: the Spearman spatial correlation between the two samples’ vectorized behavioral cross-correlation matrices was .998. Furthermore, excluding the 579 related children did not have a greater effect on the correlation between any two behavioral measures than excluding 579 randomly selected children (non-parametric *p* values > Bonferroni-corrected .05; see *Materials and methods*). In other words, family structure did not have a significant impact on the observed pattern of behavioral relationships.

### Behavioral relationships are robust to age, sex, outliers, and missing data

Control analyses confirmed that behavioral relationships were robust to potential confounds (Supplementary Fig. 1). Specifically, the overall pattern of relationships was consistent after controlling for age and sex with partial correlation (*n* = 4398; partial *rs* = .993), excluding outlier values (> 2.5 standard deviations from the group mean; *n* = 4398; *rs* = .994), excluding children with any missing behavioral scores (*n* = 2304; *rs* = .990), excluding children who completed any neuroimaging task (i.e., the emotional *n*-back, stop signal, or monetary incentive delay task; see *Materials and methods*) on a laptop outside the scanner (*n* = 3604; *rs* = .996), and excluding children with neuroimaging task performance flags provided by the ABCD study (*n* = 2154; *rs* = .974). The overall pattern of relationships was replicated to a lesser degree in a conservative subsample of children excluding relatives, outlier values, incomplete cases, children who completed neuroimaging tasks outside the scanner, and children with neuroimaging task performance flags, and controlling for age and sex (*n* = 1471; partial *rs* = .71). Finally, associations between behavioral measures were similar across data collection sites despite differences in target sociodemographics (57). Similarity between site-specific behavioral cross-correlation patterns ranged from *rs* = .42–.86 (mean *rs* = .67, *s.d.* = .10), with sites with more participants showing more typical patterns (Spearman correlation between each site’s sample size and mean similarity to all other sites = .79, *p* = 3.03×10^−5^).

### Behavioral relationships vary across the working memory spectrum

Do associations between working memory and other cognitive measures differ between children with stronger and weaker memory function? To assess this possibility, children were divided into quartiles based on their list sorting working memory performance (per-quartile *n* = 1086). Overall patterns of behavioral relationships were similar across quartiles (*rs* values > .749). Qualitatively, however, correlations between working memory and NIH Toolbox test, RAVLT, and matrix reasoning performance followed a U-shaped pattern, such that they tended to be higher in children with the weakest and strongest working memory function (see *Materials and methods* and Supplementary >Fig. 2 for quantitative comparisons).

### Neural signatures of working memory

To identify a neural signature of working memory—that is, a pattern of functional MRI activity associated with working memory function—we related out-of-scanner list sorting working memory performance to fMRI activation in response to a memory challenge. Memory-related fMRI activation was measured with a linear contrast of 2-back vs. 0-back emotional *n*-back task blocks (3444 data sets available; *n* = 2116 after exclusion; see *Materials and methods*). Subject-specific beta weights were entered into a multiple regression model including list sorting performance as a predictor with FSL’s PALM software (58). Four covariates were also included in the model: age and sex (to account for effects present in the uncorrected list sorting standard scores) (56), scanner (to account for magnet-related differences between the 26 scanners as well as effects of participant population [e.g., family income, education, race and ethnicity]), and fluid intelligence (to account for non-specific effects of cognitive function). Nonparametric significance was assessed with 1,000 permutations per contrast using PALM. Regression coefficients surviving a family-wise error-corrected *p*-value threshold of .05 were considered significant.

Working memory function was significantly related to 2-back vs. 0-back (i.e., high vs. low memory load) activation in regions of frontal and parietal cortex including bilateral intraparietal sulci, dorsal premotor cortex/frontal eye fields, dorsolateral prefrontal cortex, anterior insula, dorsal anterior cingulate cortex extending into the pre-supplementary motor area, and precuneus (Fig. 4). In line with previous work highlighting the importance of frontoparietal regions for working memory in development (32, 43, 45), children with better out-of-scanner working memory performance showed increased activity during high relative to low memory load task blocks in this distributed set of regions that overlap with frontoparietal and dorsal attention networks (59, 60) (Fig. 5).

**Figure 4.**
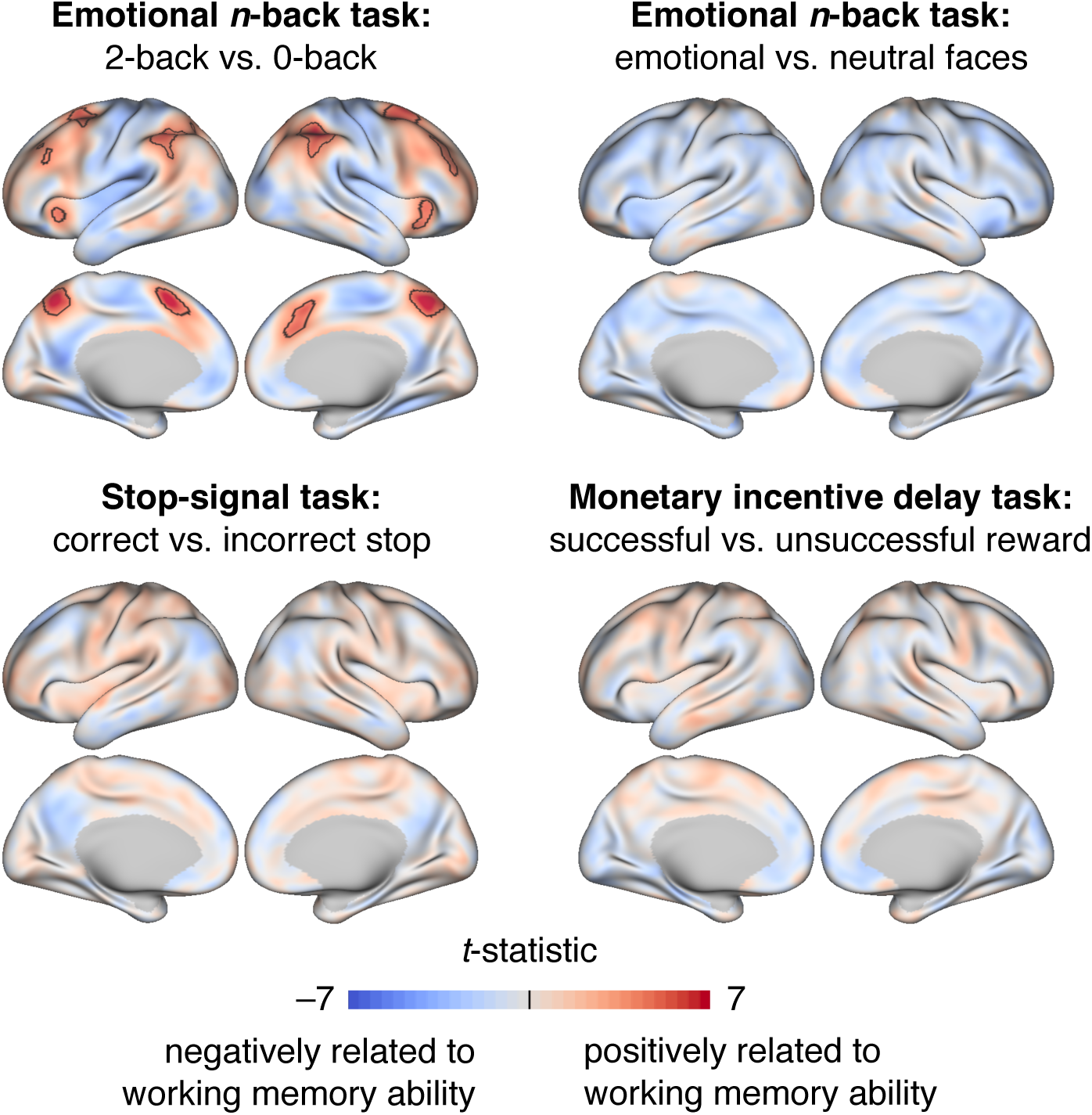
Relationships between fMRI activation and working memory function across individuals. Unthresholded *t*-statistics are visualized on the inflated cortical surface. Black outlines indicate vertices significant at family-wise error-corrected, two-tailed *p* <.05.

**Figure 5.**
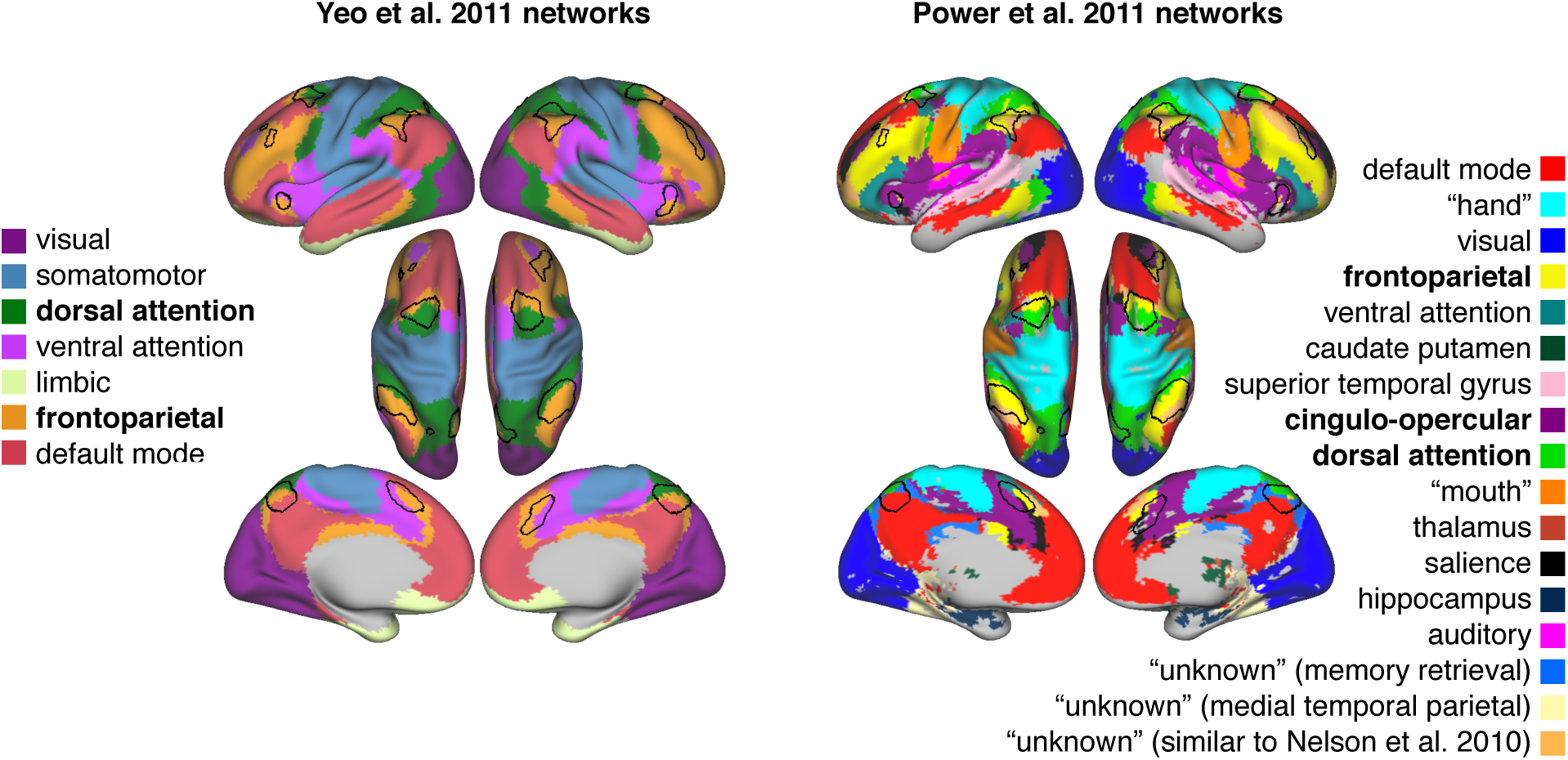
Overlap between neural signatures of working memory in childhood and canonical resting-state functional networks from Yeo et al. (2011) and Power et al. (2011). Black outlines indicate significant relationships between 2-back vs. 0-back activation and working memory function across individuals (family-wise error-corrected, two-tailed *p* < .05).

### Neural signatures of working memory are domain-specific specific

Are patterns of fMRI activation that track individual differences in working memory driven by general task demands, or are they driven by working memory engagement *per se*? We performed two analyses to disentangle these alternatives. First, we examined the association between individual differences in working memory performance and activation revealed by a contrast of emotional vs. neutral face blocks in the *n*-back task. Although these emotion-related activation patterns were measured during a working memory task, they do not reflect a working memory challenge. Therefore, significant relationships between these patterns and working memory would suggest that neural signatures of working memory are domain-general rather than domain-specific. Second, we examined the relationships between individual differences in working memory and activation patterns reflecting distinct cognitive processes in distinct task contexts: inhibitory control during a stop-signal task and reward processing during a monetary incentive delay task. Subject-specific beta coefficient maps reflecting inhibitory control-related activity were computed by contrasting successful vs. unsuccessful stop trials on the stop-signal task (3447 data sets available; *n* = 2368 after exclusion; see *Materials and methods*). Beta coefficient maps reflecting activity related to reward sensitivity were computed by contrasting successful vs. unsuccessful reward trials (i.e., volumes corresponding to positive vs. negative feedback) on the monetary incentive delay task (3543 data sets available; *n* = 2418 after exclusion). Multiple regression models including working memory performance, age, sex, scanner, and fluid intelligence were applied to predict subject-specific beta weights.

Results revealed that working memory was not significantly associated with emotion-related activation during the emotional *n*-back task, inhibitory control-related activation during the stop-signal task, or reward-related activation during the monetary incentive delay task (Fig. 4). Although we did not compare regression coefficients across conditions because participant samples were overlapping but not identical, more participants and time points were available for the stop-signal and monetary incentive delay tasks than for the emotional *n*-back task. Thus, the presence of significant effects for the working memory contrast—but not the inhibitory control or reward processing contrasts—is not attributable to sample size or amount of data per individual, and results suggest that frontoparietal activity is a domain-specific rather than a domain-general signature of working memory.

### Memory-related frontoparietal activity reflects in-scanner and out-of-scanner working memory performance

One potential explanation of the current results is that in-scanner emotional *n*-back performance—a state-like measure of working memory and task engagement rather than a measure of individual differences in working memory *per se*—drives the selective relationship between working memory and 2-back vs. 0-back frontoparietal activation. To evaluate this possibility, we replicated the analysis relating out-of-scanner working memory performance to 2-back vs. 0-back activation with age, sex, scanner, and overall *n*-back accuracy included in the model as covariates. Results revealed significant clusters in superior parietal and pre-supplementary motor areas (Supplementary Fig. 3), demonstrating that memory-related activation reflects both in-scanner and out-of-scanner working memory performance.

## Discussion

Working memory is a foundational cognitive function that changes over development and varies across individuals. Here we characterize relationships between working memory, cognitive and attentional processes, and task-related brain activity in childhood using behavioral and functional MRI data from the largest developmental neuroimaging sample to date. Behavioral analyses demonstrate that children with stronger working memory abilities perform better on a range of cognitive tasks, and revealed close relationships between working memory, performance on other memory tasks, language abilities, and fluid intelligence. Functional MRI analyses of emotional *n*-back, stop-signal, and monetary incentive delay task data provide evidence that frontoparietal activation in response to an explicit memory challenge—but not in response to task demands more generally—is a reliable marker of working memory ability. Finally, a control analysis suggests that memory-related frontoparietal activity reflects individual differences in working memory above and beyond ongoing task performance.

Positive relationships between working memory, language abilities, and fluid intelligence replicate previous work on the structure of cognition in children and adults (61–63). As expected, children with stronger working memory abilities (measured with the List Sorting Working Memory Test) also showed better performance on tests of episodic memory (Picture Sequence Memory), short-term memory (Rey Auditory Verbal Learning), and low- and high-load working memory (emotional *n*-back 0- and 2-back conditions, respectively). Correlations between these measures in the full sample of 4398 children ranged from .27–.34, suggesting that they reflect both distinct and overlapping aspects of memory function. Somewhat surprisingly given established links between working memory and processing speed (64), working memory was less closely related to performance on the Pattern Comparison Processing Speed Test than to performance on every cognitive task except the stop-signal, monetary incentive delay, and intratemporal cash choice tasks. Although the strength of the relationship between working memory and processing speed (*r*_s_ = .17) is numerically similar to previous findings with the same tasks in 8-to-12-year-olds (*r* = .26; REF 65), individual differences in working memory were more strongly related to processes including executive attention and cognitive flexibility than to processing speed in the current cohort. In addition, behavioral signatures of working memory varied as a function of working memory ability, such that relationships between behavioral measures were generally strongest in children with low working memory performance, potentially due to factors such as low attention-to-task or motivation. Finally, behavioral cross-correlation patterns were consistent after controlling for age and sex and excluding statistical outliers, incomplete cases, and neuroimaging task data collected outside the scanner. These behavioral patterns remained unchanged when measured in a subsample of the data that did not include relatives (i.e., only including one child per identified family). Thus, although it is important to account for these factors in large datasets such as the ABCD sample, the current results appear robust to effects of statistical dependence and outliers.

Neuroimaging results likewise align with previous work, providing evidence that frontoparietal activity reflects differences in working memory function during development (32, 45). The narrow age range of the current sample, however, allowed us to disentangle individual differences from developmental changes, providing novel evidence that frontoparietal brain function underlies variability in working memory both within and across individuals. Furthermore, assessing relationships between working memory and fMRI activity related to memory, emotion processing, inhibitory control, and reward processing demonstrated that frontoparietal activation is a domain-specific rather than a task-general neural signature of working memory. Accounting for in-scanner emotional *n*-back performance, which reflects individual differences in working memory and attentional processes as well as transient attentional state, revealed relationships between out-of-scanner working memory performance and memory-related fMRI activation in regions of superior parietal and pre-supplementary motor cortex. Children with stronger working memory abilities, therefore, show increased frontoparietal activation during high relative to low memory load task blocks in part because they simply perform better on these tasks, but also because of individual differences in their ability to hold and manipulate information in mind.

The current results suggest that frontoparietal activation is a domain-specific neural signature of working memory in that individual differences in working memory are selectively reflected in 2-back versus 0-back frontoparietal activity. However, frontoparietal activity does not *only* support working memory function, but is also related to processes including attention and cognitive control (66–70). Recent work has emphasized the multifunctional nature of the frontoparietal network, proposing that it represents a domain-general “cognitive core” of the brain (40). Our results are not inconsistent with this conceptualization, but demonstrate that a high versus low memory load contrast reveals a frontoparietal activity signature of working memory, and leave open the possibility that an attention or cognitive control contrast could reveal a frontoparietal activity signature of attention or cognitive control. Future work that expands the collection of attention and control tasks and varies their cognitive demands will provide additional insights into the functional significance of overlapping and distinct patterns of frontoparietal activity across psychological tasks with development.

A neural signature of working memory based on task activation data complements a growing body of work identifying neuromarkers of individual differences from functional brain connectivity. In particular, patterns of task-based and resting-state functional connectivity, or statistical dependence between two brain regions’ activity time courses, have been used to predict individual differences in abilities including attention, fluid intelligence, and aspects of memory (71–77). Recent work suggests that models based on task connectivity generally outperform those based on resting-state connectivity for predicting behavior, potentially because tasks engage circuits related to a process of interest to magnify individual differences in behaviorally relevant neural phenotypes, thereby improving predictions (55, 78–80). It is an open question, however, whether tasks selectively enhance the prediction of task-relevant behaviors. Here, motivated by previous work relating frontoparietal activation to developmental change in working memory (32, 45–47), we address this question with task activation rather than functional connectivity analyses. The current result—that frontoparietal activity indexes working memory *only when working memory is explicitly taxed*—suggests that task challenges may reveal neural signatures of task-relevant behaviors, and underscores the importance of multi-task or multi-condition data for elucidating state-specific and state-general biomarkers of behavior.

The goal of the current work was to characterize a brain-based biomarker and behavioral signature of working memory in childhood not just for the sake of understanding these relationships at a single point in time, but also for understanding their trajectories across development. Because the ABCD study will follow children from age 9–10 to age 19–20, longitudinal work can provide new insights into associations between working memory, cognitive and attentional processes, and real-world outcomes across adolescence and young adulthood. Biennial MRI sessions— during which participants will complete the same emotional *n*-back, stop-signal, and monetary incentive delay tasks that they completed at age nine and ten—will also facilitate the discovery of changing neural signatures of abilities and behavior. For example, will there be changes in the distinct and overlapping brain activity patterns associated with working memory, inhibitory control, and reward processing with age? Will the domain-specificity and domain-generality of these signatures vary over time? Are there different developmental trajectories for frontoparietal organization of function across these processes? A fruitful way to frame the current findings is as a single point along a nonlinear trajectory rather than as a summary of working memory function in development as a whole.

Finally, as sample sizes in psychology and human neuroscience rapidly increase, it is important to note limitations of big data cohort-based approaches. First, behavioral and neuroimaging task batteries for these studies are determined by committee to address specific scientific goals. Although the resulting task sets often assess a range of mental processes, they may not be optimal for answering all questions. In the ABCD study neuroimaging battery, for example, cognitive control demands and task difficulty are not equated across the emotional *n*-back, stop-signal, and monetary incentive delay tasks. Thus, the 2-back vs. 0-back contrast may reflect processes such as cognitive control and attention that are not reflected in the three control contrasts. Future work relating individual differences in working memory to fMRI activity reflecting cognitive control, attentional engagement, and other processes in contexts matched for task difficulty will further inform the domain-specificity and -generality of neural signatures of working memory. Second, large samples are not necessarily representative samples, and the ABCD cohort, while geographically, demographically, and socioeconomically diverse, should not be considered representative of the country or world as a whole (57). Looking ahead, future work relating cognitive and neural measures in weighted samples (81) can complement existing studies of single- and multi-site datasets. Third, just as the ABCD participant population may not represent youth as a whole, the structure of neurocognition in nine- and ten-year-olds likely does not reflect that of children at other ages. Longitudinal analyses of the ABCD cohort can inform changes in brain–behavior relationships across adolescence, and data collection efforts such as the Human Connectome Project (HCP) Development Study (82) and HCP Aging study (83) can inform these associations in younger and older individuals. Finally, because even small effects can reach significance when samples are large, it is helpful to distinguish statistical from practical significance. Here we focused on statistical significance as a proof-of-principle demonstration that memory-related frontoparietal activity tracks individual differences in working memory in childhood. Future work can evaluate practical or applied significance by testing whether models based on task activation patterns generalize to predict real-world outcomes including academic performance or changes in these outcomes over time.

Despite these limitations, the current results provide the most well powered characterization of relationships between working memory, cognitive and attentional processes, and task-based fMRI activation in development to date. Overall, they replicate established findings that children with stronger working memory function perform better on a variety of cognitive tasks, particularly those assessing other aspects of memory, language skills, and fluid intelligence. Furthermore, they provide evidence that frontoparietal network activation in response to an explicit memory challenge is a robust and domain-specific marker of individual differences in working memory ability at age nine and ten. Together these results inform understanding of the structure of neurocognition in childhood, and highlight the importance of evaluating brain– behavior relationships in multiple task contexts to demarcate the specificity and generality of neural signatures of abilities and behavior.

## Materials and methods

### The Adolescent Brain Cognitive Development (ABCD) study sample

The ABCD study is a multi-site study following a geographically, demographically, and socioeconomically diverse sample of over 11,000 children in the United States from age 9–10 to age 19–20. Launched in September 2016, the study aims to characterize cognitive and neural development with measures of neurocognition, physical and mental health, social and emotional function, and culture and environment. Exclusionary criteria include a diagnosis of schizophrenia, a moderate to severe autism spectrum disorder, an intellectual disability, or a substance use disorder at recruitment. Children with a persistent major neurological disorder (e.g., cerebral palsy, a brain tumor, stroke, brain aneurysm, brain hemorrhage, subdural hematoma), multiple sclerosis, sickle cell disease, or certain seizure disorders (Lennox-Gastaut syndrome, Dravet syndrome, and Landau Kleffner syndrome) were also excluded. Data collection includes yearly behavioral assessments, interviews, questionnaires, and biosample collection as well as biennial MRI scans (54). Here we analyze year-one (baseline) demographic and behavioral data from the first half of the cohort (*n* = 4521; 47.5% female), collected across 21 sites when children were 9–10 years old and made available as part of curated data release 1.1 on October 15, 2018 (DOI 10.15154/1412097). Sample demographics including race, ethnicity, socioeconomic status, and symptoms of internalizing and externalizing disorders are available in REF 61.

### Behavioral data and exclusion criteria

To characterize associations between working memory, measured with the NIH Toolbox List Sorting Working Memory Test, and other cognitive abilities in childhood, we analyzed performance data from all available neurocognitive (56) and neuroimaging (54) tasks. Data from children diagnosed with autism spectrum disorder or epilepsy were excluded from analysis as moderate to severe forms of autism spectrum disorder and other seizure disorders were exclusionary for the study (*n* = 123). Data from children with attention deficit hyperactivity disorder, depression, bipolar disorder, anxiety, and phobias (*n* = 618) were not excluded, as these diagnoses were assessed with a single screening question and we aimed to characterize working memory in a heterogeneous developmental population. Task performance measures, described here and in Supplementary Table 1, were selected based on previous work including reports of ABCD baseline data (54, 56).

### NIH Toolbox cognition battery

The NIH Toolbox^®^ cognition battery includes seven tasks measuring multiple aspects of cognition (84) (Supplementary Table 1, column 3). Performance is measured using uncorrected standard scores as age-corrected scores are currently undergoing revision by the NIH Toolbox (56).

#### The Toolbox List Sorting Working Memory Test ^®^

measures working memory by asking children to recall stimuli in different orders (2). Children are shown pictures and hear the names of animals and foods of different sizes. They are then asked to repeat back the items in order from smallest to largest. Children are first shown pictures of two animals, and then shown longer lists (up to seven) if they respond correctly. Children are next shown pictures of animals and foods together, and are asked to repeat the animals in order of size and then the foods in order of size. Interleaved lists increase in length from two to seven if children respond accurately. Performance scores reflect the number of accurate responses.

#### The Toolbox Picture Vocabulary Test ^®^

measures language and verbal abilities (85). Children hear a series of words, and are asked to choose which of four pictures most closely matches the meaning of the word.

#### The Toolbox Flanker Task ^®^

was adapted from the Attention Network Task (86), a flanker task (87) used to measure cognitive control and attention. On each trial, children see a row of five arrows. The outer four arrows (distractors, or flankers) all point to the left or right of the screen. The middle arrow (the target) points in the same direction as the flankers on congruent trials, and the opposite direction of the flankers on incongruent trials. Children are asked to indicate whether the center arrow points to the left or to the right. Performance scores are based on speed and accuracy.

#### The Toolbox Dimensional Change Card Sort Task ^®^

measures cognitive flexibility (88). On each trial, children see two objects on a screen. They are asked to match a third item with one of the initial two based on either color or shape. Children first match items based on one dimension (e.g., color), then match items based on the other dimension (e.g., shape), and finally match based on both shape and color in pseudorandom order. Performance scores are based on speed and accuracy.

#### The Toolbox Pattern Comparison Processing Speed Test ^®^

measures visual processing speed (65, 89). Children are shown two pictures and are asked to indicate whether they are the same or different. Scores are based on the number of correct responses within a time limit.

#### The Toolbox Picture Sequence Memory Test ^®^

measures episodic memory and visuospatial sequencing (90). Children are shown 15 pictures of activities or events and asked to reproduce the presentation order.

#### The Toolbox Oral Reading Recognition Task ^®^

measures reading abilities by asking children to pronounce a series of written letters and words (85).

### Matrix reasoning

The matrix reasoning subtest of the Wechsler Intelligence Test for Children-V (WISC-V) (91) measures fluid and spatial reasoning, perceptual organization, visual attention, and sequencing. On each trial, children are shown an array of visual stimuli, and are asked to select one of four stimuli that best completes the pattern. The task continues until a child makes three consecutive errors or completes 32 trials. Performance is measured by converting the number of total correct items to a standard score (56).

### Rey Auditory Verbal Learning

The Rey Auditory Verbal Learning Test (RAVLT) measures learning and memory. During the test, children hear a list of 15 unrelated words five times. Each time they hear the list, they are asked to recall as many words as possible. After these five learning trials, children hear a distractor list and are again asked to recall as many words as they can. Recall of the initial list is assessed immediately after the distractor list and again 30 minutes later (56). Here we measure performance as the number of correctly recalled words on these immediate and delayed memory assessments (i.e., RAVLT trials vi and vii).

### Intertemporal cash choice

The intertemporal cash choice task (92) assesses children’s delay of gratification, motivation, and impulsivity (56). Children are asked, “Let’s pretend a kind person wanted to give you some money. Would you rather have $75 in three days or $115 in 3 months?” Smaller-sooner reward choices were coded with a “1”, larger-later reward choices were coded with a “2”, and infrequent “don’t know” responses were excluded from analysis.

### Little Man

The Little Man task (93) measures aspects of visuospatial processing including mental rotation. During this task, children see a cartoon of a man holding a briefcase in his left or right hand appear on a computer screen. The man can be right side up or upside down, and can appear facing the child or with his back turned. Children are asked to indicate whether the man’s briefcase is in his left or right hand via button press. The task includes practice trials and 32 assessment trials. Performance is measured with efficiency (percent accuracy divided by mean correct-trial response time) (56).

### Emotional *n*-back

The in-scanner emotional *n*-back (EN-back) task engages processes related to memory and emotion regulation (54, 94). During the task, children perform 0-back (low memory load) and 2-back (high memory load) task blocks with four types of stimuli: happy, fearful, and neutral face photographs (95, 96) and place photographs. Data are collected during two approximately 5-min functional MRI runs each with four 0-back and 2-back blocks each. Runs included 362 whole-brain volumes after discarded acquisitions. At the start of each 0-back block, children are shown a target stimulus and asked to press a button corresponding to “match” when they see an identical picture and a button corresponding to “no match” when they see a different picture. During 2-back blocks, children are asked to press “match” when they see a picture identical to the one they saw two trials back. Performance is quantified as percent accuracy on 0-back and 2-back blocks.

### Recognition memory

After scanning, memory for EN-back task stimuli is assessed with a recognition memory test (54, 94). During this test, children are presented with 48 EN-back stimuli and 48 novel stimuli (i.e., 12 old and new happy, fearful, and neutral face photographs and 12 old and new places), and are asked to rate whether each picture is “old” or “new.” Performance is assessed with sensitivity (*d’*) averaged across stimulus types.

### Stop-signal

The in-scanner stop-signal task (97) (SST) is designed to measure impulsivity and impulse control (54). SST data are collected during two approximately 6-minute functional MRI runs (437 volumes after discarded acquisitions) each with 180 trials each. On each trial, children see an arrow pointing to the left or to the right of the screen (the go signal). They are instructed to indicate the direction of the arrow with a button press as quickly and accurately as possible, except when an upright arrow (the stop signal) appears on the screen (16.67% of trials). The time between go and stop signal onset, the stop-signal delay, is staircased so that each child achieves approximately 50% accuracy on stop trials. Performance is measured with stop-signal reaction time (SSRT), or the mean stop-signal delay subtracted from the mean reaction time on correct go trials. For consistency with other behavioral measures, SSRTs were reverse scored (multiplied by –1) so that higher scores correspond to better performance.

### Monetary incentive delay

The in-scanner monetary incentive delay task (98, 99) (MID) measures aspects of reward processing, including anticipation and receipt of rewards and losses and motivation to earn rewards and avoid losses (54). Data are collected during two approximately 5.5-minute, 50-trial functional MRI runs (403 volumes per run after discarded acquisitions). Trials begin with a cue indicating whether the child can win $.20 or $5, lose $.20 or $5, or earn $0. After 1500–4000 ms, a target appears for 150–500 ms. Target timing is staircased such that each child achieves approximately 60% accuracy. Children must respond during the target presentation to achieve the indicated trial outcome. Trials are followed by feedback indicating the outcome. Overall task performance is summarized as the average amount of money earned during both runs.

### Relationships between behavioral measures

To establish whether associations between working memory and other cognitive abilities were robust to potential confounds such as age, sex, missing data, outliers, and statistical dependence introduced by family structure, data collection method, and site, we first cross-correlated behavioral measures using data from all children meeting inclusion criteria (*n* = 4398). Although normality was not evaluated with formal tests, which reject the null hypothesis for near-normal distributions in large samples (100), rank correlation was applied because visual inspection indicated that behavioral measures were not normally distributed (Fig. 1). We subsequently replicated this analysis using:

1. data from only one child per family based on self-report (*n* = 3819). For families with multiple children in the 4398-participant cohort, the child whose randomly assigned NDAR Global Unique Identifier (GUID) came first in alphabetical order was included in this sample.
2. data from all children meeting inclusion criteria using Spearman partial correlation to control for age and sex
3. data values within 2.5 standard deviations of the group mean (see Supplementary Table 1)
4. data from children with no missing values (i.e., complete cases; *n* = 2304)
5. data from children who completed the emotional *n*-back, SST, and MID tasks during MRI data collection rather than on a laptop outside the scanner (*n* = 3604)
6. data from children without performance flags on the emotional *n*-back, SST, and MID tasks (*n* = 2154). Performance flags, provided in ABCD Release 1.1, were assigned based on the following criteria: <60% 0-back or 2-back accuracy on the emotional *n*-back task; <150 go trials, <60% go trial accuracy, >30% incorrect go trial percentage, >30% late go trial percentage, >30% “no response” go trials, <30 stop trials, or <20% or >80% stop trial accuracy on the SST; <3 positive and negative feedback events for large reward, small reward, large loss, small loss, or no stakes trials on the MID task.
7. data from a conservative subsample excluding outlier values, incomplete cases, children with who completed neuroimaging tasks outside of the scanner, children with neuroimaging task performance flags, and relatives, and controlling for age and sex with partial correlation
8. data from each of the 20 data collection sites with more than 100 participants separately (*n* = 108–417; mean *n* = 218.15; *s.d.* = 86.86; 21^st^ site with 35 children excluded)

Due to the frequency of missing data, relationships between behavioral measures were evaluated with pairwise correlations rather than with data reduction techniques such as principal component analysis (PCA), which do not typically allow for missing data. 47.61% of children were missing at least one performance measure, and neuroimaging task performance data were missing in 27.95% of the sample on average (Supplementary Table 1; although note that recovery of missing data is ongoing). Although Bayesian probabilistic PCA can account for missing data as well as the nesting of participants in families and data collection sites (61), this approach assumes that missing data occur randomly, independent of other sample features (101). This assumption is violated in the current sample, as, for example, children with better working memory function are less likely to be missing other behavioral measures (Spearman correlation between list sorting performance and number of missing performance measures = –0.06, *p* = 6.39×10^−5^).

### Effects of family structure on behavioral relationships

Given the importance of accounting for family structure in the ABCD sample, we characterized effects of relatedness on behavioral relationships with an additional analysis. First, we computed the absolute difference between all 136 pairwise behavioral correlations in the full sample (*n* = 4398) and the unrelated subsample (*n* = 3819 after excluding 579 related children). Next, we randomly excluded 579 children from the full sample, re-calculated pairwise behavioral relationships, and recorded the difference between the full-sample correlations and these random subsample correlations. We repeated this process 1,000 times to generate a null distribution of correlation coefficient differences for each pair of behavioral measures. Non-parametric *p*-values were generated by comparing each true correlation difference, |*r*(*i,j*)_full sample_ – *r*(*i,j*)_unrelated subsample_|, to its corresponding null distribution. We elected to take this conservative sub-sampling approach rather than to control for family relatedness with linear mixed models given the complexity of possible relationships (e.g., monozygotic and dizygotic twins, full siblings, half siblings, cousins, etc.) and the fact that relatedness may be inaccurately captured with self-report measures.

Using the subsampling approach, we found that 133 of the 136 pairwise behavioral relationships did not differ between the full sample and the unrelated subsample more than they differed between the full sample and the random subsamples (uncorrected non-parametric *p* value range = .056 – .99). Excluding relatives had a larger effect (uncorrected *p* < .05) than excluding random participants on three correlations: mean monetary incentive delay (MID) earnings vs. emotional *n*-back recognition memory *d’* (*p* = .011), 2-back accuracy vs. NIH Toolbox Picture Vocabulary (*p* = .028), and 2-back accuracy vs. NIH Toolbox Picture Vocabulary (*p* = .033). None of these relationships, however, survive Bonferroni correction for 136 comparisons (*p* = .05/136 = 3.68×10^−4^). Thus, excluding family members from the sample did not disproportionately affect pairwise behavioral relationships.

### Behavioral relationships across the working memory spectrum

Working memory may be differentially associated with other cognitive abilities in children with better and worse working memory function. To assess this possibility, we recomputed behavioral relationships in each quartile (defined using list sorting performance) of the 4398-child sample and measured the similarity of patterns of behavioral relationships with spatial correlation between all pairs of quartiles. Quartile-specific patterns are visualized in Supplementary Fig. 2.

We next compared relationships between working memory and other abilities across all pairs of quartiles. This analysis revealed significantly stronger relationships between working memory and picture sequence memory, Rey Auditory Verbal Learning (RAVLT) immediate and delayed recall, and matrix reasoning in the lowest (i.e., first) quartile than the third quartile (*z* values > 3.95; *p* < 7.90×10^−5^). The relationship between working memory and RAVLT delayed recall was also stronger in the first than second quartile (*z* = 3.53, *p* = 4.10×10^−4^). Correlations between working memory and these other memory measures (picture sequence memory and RAVLT immediate and delayed recall) did not continue to decrease in children with the strongest working memory function, but rather were significantly stronger in the fourth than third quartile (*z* values > 4.39; *p* < 1.13×10^−5^). Finally, the correlation between working memory and cash choice task selection was higher in the third than first quartile (*z* = 3.77, *p* = 1.66×10^−4^). No other pairwise comparisons reached statistical significance.

### Neuroimaging data collection

ABCD scan sessions included a localizer and acquisition of a high-resolution anatomical scan, two runs of resting state fMRI, diffusion weighted images, 3D T2-weighted spin echo images, two more runs of resting state fMRI, and task-based fMRI. (Sites with Siemens scanners used Framewise Integrated Real-time MRI Monitoring (102) [FIRMM] to monitor children’s head motion during data collection. Scan operators at these sites may have stopped resting-state data collection after three runs if 12.5 minutes of low-motion resting-state data had been collected.) Image acquisition order was fixed, but fMRI task order was randomized across participants (54). Data were collected on Siemens Prisma, Phillips and GE 750 3T scanners, with detailed acquisition parameters reported in previous work (54, 103). Functional images were collected using a multiband gradient EPI sequence with the following parameters: TR = 800 ms, TE = 30 ms, flip angle = 52°, 60 slices acquired in the axial plane, voxel size = 2.4 mm^3^, multiband slice acceleration factor = 6.

### Image processing

Task-based data were processed by the ABCD study Data Analysis and Informatics Center (DAIC) using approaches described in detail in REF 103. Preprocessing steps included motion correction with 3dvolreg in AFNI, B_0_-distortion (i.e., field-map) correction, gradient nonlinearity distortion correction, and resampling scans into alignment with cubic interpolation using a mid-session scan as the reference. Registration between T2-weighted spin echo scans, field maps, and T1-weighted structural images was performed using mutual information. Functional images were aligned to T1-weighted images using rigid-body transformation (103).

After preprocessing, the equivalent of 16 volumes was removed from the start of each run. For Siemens and Philips scanners, 8 volumes were removed because the first 8 volumes were used as the multiband reference scans. For GE scanners running DV25 software (nearly all GE-scanner datasets included in ABCD Data Release 1.1), 5 volumes were removed because the first 12 volumes were used as the multiband reference. The images were then combined into a single volume and saved as the initial TR (leaving a total of 5 frames to be discarded). For GE scanners running DV26 software, 16 volumes were removed.

Voxel-wise time series data were next normalized with run. Task-related activity estimates were generated for each child using general linear models (GLMs) with 3dDeconvolve in AFNI (103). GLMs included nuisance regressors accounting for baseline and quadratic trends as well as motion estimates and their derivatives temporally filtered to attenuate .31–.43 Hz signals related to respiration (104). Volumes with framewise displacement values >.9 mm were censored (105).

In addition to fixation, the emotional *n*-back task GLM included predictors for happy, fearful, and neutral face as well as place stimuli in the 0-back and 2-back conditions. Task bocks (approximately 24 s) were modeled as square waves convolved with a two-parameter gamma basis function (103). The stop-signal task model included predictors for correct and incorrect stop and go trials, modeled as instantaneous. The monetary incentive delay model included small and large reward and loss cues and feedback and no stakes cues, modeled as instantaneous (103). Linear contrasts of interest for each task are described in the *Results*. GLM beta coefficients for cortical gray matter voxels were sampled into surface space. (This step differs from the processing pipeline described in REF 103, in which preprocessed data were sampled onto the cortical surface, but does not affect the beta values.)

### Neuroimaging data exclusion

Neuroimaging data from children with poor structural scan quality, determined with curated data release 1.1 sheet freesqc01.txt variable *fsqc_qc*, were excluded from analysis. For each task contrast, participants with fewer than 550 degrees of freedom in preprocessed, concatenated fMRI time series, missing grayordinate (i.e., gray-matter vertex or voxel) values, and/or or extreme values (>3 standard deviations from the group mean) for the mean or standard deviations of beta weights over all grayordinates were also excluded. All fMRI analyses were performed using data from only one child per family to avoid confounds introduced by family structure. We elected to take this conservative approach rather than attempting to control for family structure with multi-level block permutation (106) because relatedness was determined with self-report rather than with genetic testing, the gold standard.

## Data source

The ABCD behavioral, demographic, and quality control data used in this report came from NIMH Data Archive Digital Object Identifier (DOI) 10.15154/1412097. DOIs can be found at ndar.nih.gov/study.html?id=576.

## References

1. Johnson MK, et al. (2013) The relationship between working memory capacity and broad measures of cognitive ability in healthy adults and people with schizophrenia. Neuropsychology 27(2):220–229.

2. Tulsky DS, et al. (2014) NIH Toolbox Cognition Battery (NIHTB-CB): List sorting test to measure working memory. J Int Neuropsychol Soc 20(6):599–610.

3. Xu Z, Adam KCS, Fang X, Vogel EK (2018) The reliability and stability of visual working memory capacity. Behav Res Methods 50(2):576–588.

4. Alp IE (1994) Measuring the size of working memory in very young children: the imitation sorting task. Int J Behav Dev 17(1):125–141.

5. Ross RG, Wagner B, Heinlein S, Zerbe GO (2008) The stability of inhibitory and working memory deficits in children and adolescents who are children of parents with schizophrenia. Schizophr Bull 34(1):47–51.

6. Engel de Abreu PMJ, Conway ARA, Gathercole SE (2010) Working memory and fluid intelligence in young children. Intelligence 38(6):552–561.

7. Fukuda K, Vogel E, Mayr U, Awh E (2010) Quantity not quality: The relationship between fluid intelligence and working memory capacity. Psychon Bull Rev 17(5):673–679.

8. Kane MJ, Engle RW (2002) The role of prefrontal cortex in working-memory capacity, executive attention, and general fluid intelligence: An individual-differences perspective. Psychon Bull Rev 9(4):637–671.

9. Huang L, Mo L, Li Y (2012) Measuring the interrelations among multiple paradigms of visual attention: an individual differences approach. J Exp Psychol Hum Percept Perform 38(2):414–28.

10. Alloway TP, Gathercole SE, Pickering SJ (2006) Verbal and visuospatial shortterm and working memory in children: are they separable? Child Dev 77(6):1698–1716.

11. Davidson MC, Amso D, Anderson LC, Diamond A (2006) Development of cognitive control and executive functions from 4 to 13 years: Evidence from manipulations of memory, inhibition, and task switching. Neuropsychologia 44(11):2037–2078.

12. De Smedt B, et al. (2009) Working memory and individual differences in mathematics achievement: A longitudinal study from first grade to second grade. J Exp Child Psychol 103(2):186–201.

13. Bayliss DM, Jarrold C, Gunn DM, Baddeley AD (2003) The complexities of complex span: explaining individual differences in working memory in children and adults. J Exp Psychol Gen 132(1):71–92.

14. Nouwens S, Groen MA, Verhoeven L (2017) How working memory relates to children’s reading comprehension: the importance of domain-specificity in storage and processing. Read Writ 30(1):105–120.

15. Alloway TP, Alloway RG (2010) Investigating the predictive roles of working memory and IQ in academic attainment. J Exp Child Psychol 106(1):20–29.

16. Ross-Sheehy S, Oakes LM, Luck SJ (2003) The development of visual short-term memory capacity in infants. Child Dev 74(6):1807–1822.

17. Buss AT, Ross-Sheehy S, Reynolds GD (2018) Visual working memory in early development: a developmental cognitive neuroscience perspective. J Neurophysiol 120(4):1472–1483.

18. Reynolds GD, Romano AC (2016) The development of attention systems and working memory in infancy. Front Syst Neurosci 10:15.

19. Diamond A, Goldman-Rakic PS (1989) Comparison of human infants and rhesus monkeys on Piaget’s AB task: evidence for dependence on dorsolateral prefrontal cortex. Exp Brain Res 74(1):24–40.

20. Gathercole SE, Pickering SJ, Ambridge B, Wearing H (2004) The structure of working memory from 4 to 15 years of age. Dev Psychol 40(2):177–190.

21. Ullman H, Almeida R, Klingberg T (2014) Structural maturation and brain activity predict future working memory capacity during childhood development. J Neurosci 34(5):1592–1598.

22. Luciana M, Conklin HM, Hooper CJ, Yarger RS (2005) The development of nonverbal working memory and executive control processes in adolescents. Child Dev 76(3):697–712.

23. Conklin HM, Luciana M, Hooper CJ, Yarger RS (2007) Working memory performance in typically developing children and adolescents: Behavioral evidence of protracted frontal lobe development. Dev Neuropsychol 31(1):103–128.

24. Isbell E, Fukuda K, Neville HJ, Vogel EK (2015) Visual working memory continues to develop through adolescence. Front Psychol 6:696.

25. Nyberg L, et al. (2013) Age-related and genetic modulation of frontal cortex efficiency. J Cogn Neurosci 26(4):746–754.

26. Eriksson J, Vogel EK, Lansner A, Bergström F, Nyberg L (2015) Neurocognitive Architecture of Working Memory. Neuron 88(1):33–46.

27. Swanson HL (2017) Verbal and visual-spatial working memory: What develops over a life span? Dev Psychol 53(5):971–995.

28. Alloway TP, Alloway RG (2013) Working memory across the lifespan: A crosssectional approach. J Cogn Psychol 25(1):84–93.

29. Fry AF, Hale S (1996) Processing speed, working memory, and fluid intelligence: evidence for a developmental cascade. Psychol Sci 7(4):237–241.

30. Fry AF, Hale S (2000) Relationships among processing speed, working memory, and fluid intelligence in children. Biol Psychol 54(1):1–34.

31. Pailian H, Libertus ME, Feigenson L, Halberda J (2016) Visual working memory capacity increases between ages 3 and 8 years, controlling for gains in attention, perception, and executive control. Attention, Perception, Psychophys 78(6):1556–1573.

32. Klingberg T, Forssberg H, Westerberg H (2002) Increased brain activity in frontal and parietal cortex underlies the development of visuospatial working memory capacity during childhood. J Cogn Neurosci 14(1):1–10.

33. Casey BJ, Giedd JN, Thomas KM (2000) Structural and functional brain development and its relation to cognitive development. Biol Psychol 54(1):241–257.

34. Casey BJ, Tottenham N, Liston C, Durston S (2005) Imaging the developing brain: what have we learned about cognitive development? Trends Cogn Sci 9(3):104–110.

35. Steinberg L, Cauffman E, Woolard J, Graham S, Banich M (2009) Are adolescents less mature than adults?: Minors’ access to abortion, the juvenile death penalty, and the alleged APA “flip-flop.” Am Psychol 64(7):583–594.

36. Salthouse TA, Babcock RL (1991) Decomposing adult age differences in working memory. Dev Psychol 27(5):763–776.

37. McNab F, et al. (2015) Age-related changes in working memory and the ability to ignore distraction. Proc Natl Acad Sci 112(20):6515–6518.

38. Gazzaley A, Cooney JW, Rissman J, D’Esposito M (2005) Top-down suppression deficit underlies working memory impairment in normal aging. Nat Neurosci 8(10):1298–1300.

39. Woolgar A, Hampshire A, Thompson R, Duncan J (2011) Adaptive coding of task-relevant information in human frontoparietal cortex. J Neurosci 31(41):14592–14599.

40. Assem M, Glasser MF, Van Essen DC, Duncan J (2019) A domain-general cognitive core defined in multimodally parcellated human cortex. bioRxiv:517599.

41. Scolari M, Seidl-Rathkopf KN, Kastner S (2015) Functions of the human frontoparietal attention network: Evidence from neuroimaging. Curr Opin Behav Sci 1:32–39.

42. Casey BJ, et al. (1995) Activation of prefrontal cortex in children during a nonspatial working memory task with functional mri. Neuroimage.

43. Klingberg T, Darki F (2014) The role of fronto-parietal and fronto-striatal networks in the development of working memory: a longitudinal study. Cereb Cortex 25(6):1587–1595.

44. Tamnes CK, et al. (2013) Longitudinal working memory development is related to structural maturation of frontal and parietal cortices. J Cogn Neurosci 25(10):1611–1623.

45. Satterthwaite TD, et al. (2013) Functional maturation of the executive system during adolescence. J Neurosci 33(41):16249–16261.

46. Kwon H, Reiss AL, Menon V (2002) Neural basis of protracted developmental changes in visuo-spatial working memory. Proc Natl Acad Sci 99(20):13336–13341.

47. Crone EA, Wendelken C, van Leijenhorst L, Donohue S, Bunge SA (2006) Neurocognitive development of the ability to manipulate information in working memory. Proc Natl Acad Sci 103(24):9315–9320.

48. Tittgemeyer M, Fiebach CJ, Ekman M, Melzer C, Derrfuss J (2016) Different roles of direct and indirect frontoparietal pathways for individual working memory capacity. J Neurosci 36(10):2894–2903.

49. Burzynska AZ, et al. (2011) Microstructure of frontoparietal connections predicts cortical responsivity and working memory performance. Cereb Cortex 21(10):2261–2271.

50. Takeuchi H, et al. (2011) Verbal working memory performance correlates with regional white matter structures in the frontoparietal regions. Neuropsychologia 49(12):3466–3473.

51. Palva JM, Monto S, Kulashekhar S, Palva S (2010) Neuronal synchrony reveals working memory networks and predicts individual memory capacity. Proc Natl Acad Sci 107(16):7580–7585.

52. Osaka M, et al. (2003) The neural basis of individual differences in working memory capacity: an fMRI study. Neuroimage 18(3):789–797.

53. Volkow ND, et al. (2018) The conception of the ABCD study: From substance use to a broad NIH collaboration. Dev Cogn Neurosci 32:4–7.

54. Casey BJ, et al. (2018) The Adolescent Brain Cognitive Development (ABCD) study: Imaging acquisition across 21 sites. Dev Cogn Neurosci 32:43–54.

55. Finn ES, et al. (2017) Can brain state be manipulated to emphasize individual differences in functional connectivity? Neuroimage 160:140–151.

56. Luciana M, et al. (2018) Adolescent neurocognitive development and impacts of substance use: Overview of the adolescent brain cognitive development (ABCD) baseline neurocognition battery. Dev Cogn Neurosci 32:67–79.

57. Garavan H, et al. (2018) Recruiting the ABCD sample: Design considerations and procedures. Dev Cogn Neurosci 32:16–22.

58. Winkler AM, Ridgway GR, Webster MA, Smith SM, Nichols TE (2014) Permutation inference for the general linear model. Neuroimage 92:381–397.

59. Power JD, et al. (2011) Functional network organization of the human brain. Neuron 72(4):665–678.

60. Yeo BTT, et al. (2011) The organization of the human cerebral cortex estimated by intrinsic functional connectivity. J Neurophysiol 106(3):1125–65.

61. Thompson WK, et al. (2018) The structure of cognition in 9 and 10 year-old children and associations with problem behaviors: Findings from the ABCD study’s baseline neurocognitive battery. Dev Cogn Neurosci 36: 100606.

62. Engle RW, Tuholski SW, Laughlin JE, Conway ARA (1999) Working memory, short-term memory, and general fluid intelligence: A latent-variable approach. J Exp Psychol Gen 128(3):309–331.

63. Gathercole SE (1999) Cognitive approaches to the development of short-term memory. Trends Cogn Sci 3(11):410–419.

64. Conway ARA, Cowan N, Bunting MF, Therriault DJ, Minkoff SRB (2002) A latent variable analysis of working memory capacity, short-term memory capacity, processing speed, and general fluid intelligence. Intelligence 30(2):163–183.

65. Carlozzi NE, Beaumont JL, Tulsky DS, Gershon RC (2015) The NIH toolbox pattern comparison processing speed test: normative data. Arch Clin Neuropsychol 30(5):359–368.

66. Corbetta M, Shulman GL (2002) Control of goal-directed and stimulus-driven attention in the brain. Nat Rev Neurosci 3:201–215.

67. Vincent JL, Kahn I, Snyder AZ, Raichle ME, Buckner RL (2008) Evidence for a frontoparietal control system revealed by intrinsic functional connectivity. J Neurophysiol 100(6):3328–3342.

68. Spreng RN, Stevens WD, Chamberlain JP, Gilmore AW, Schacter DL (2010) Default network activity, coupled with the frontoparietal control network, supports goal-directed cognition. Neuroimage 53(1):303–317.

69. Spreng RN, Sepulcre J, Turner GR, Stevens WD, Schacter DL (2012) Intrinsic architecture underlying the relations among the default, dorsal attention, and fronto-parietal control networks of the human brain. J Cogn Neurosci:1–12.

70. Ptak R (2011) The frontoparietal attention network of the human brain: action, saliency, and a priority map of the environment. Neurosci 18(5):502–515.

71. Finn ES, et al. (2015) Functional connectome fingerprinting: identifying individuals using patterns of brain connectivity. Nat Neurosci 18(11):1664–1671.

72. Rosenberg MD, Finn ES, Scheinost D, Constable RT, Chun MM (2017) Characterizing attention with predictive network models. Trends Cogn Sci 21(4):290–302.

73. Lin Q, et al. (2018) Resting-state functional connectivity predicts cognitive impairment related to Alzheimer’s disease. Front Aging Neurosci 10.

74. Avery EW, et al. Whole-brain functional connectivity predicts working memory performance in novel healthy and memory-impaired individuals. Program No. 426.16. 2018 Neuroscience Meeting Planner. San Diego, CA: Society for Neuroscience, 2018. Online. (2018).

75. Yamashita M, et al. (2018) A prediction model of working memory across health and psychiatric disease using whole-brain functional connectivity. Elife 7:e38844.

76. Galeano Weber EM, Hahn T, Hilger K, Fiebach CJ (2017) Distributed patterns of occipito-parietal functional connectivity predict the precision of visual working memory. Neuroimage 146:404–418.

77. Rudolph MD, et al. (2018) Maternal IL-6 during pregnancy can be estimated from the newborn brain connectome and predicts future working memory performance in offspring. Nat Neurosci 21(5):765–772.

78. Greene AS, Gao S, Scheinost D, Constable RT (2018) Task-induced brain state manipulation improves prediction of individual traits. Nat Commun 9(1):2807.

79. Yoo K, et al. (2018) Connectome-based predictive modeling of attention: Comparing different functional connectivity features and prediction methods across datasets. Neuroimage 167:11–22.

80. Rosenberg MD, Casey BJ, Holmes AJ (2018) Prediction complements explanation in understanding the developing brain. Nat Commun 9(1):589.

81. LeWinn KZ, Sheridan MA, Keyes KM, Hamilton A, McLaughlin KA (2017) Sample composition alters associations between age and brain structure. Nat Commun 8(1):874.

82. Somerville LH, et al. (2018) The Lifespan Human Connectome Project in Development: A large-scale study of brain connectivity development in 5–21 year olds. Neuroimage 183:456–468.

83. Bookheimer SY, et al. (2019) The Lifespan Human Connectome Project in Aging: An overview. Neuroimage 185:335–348.

84. Gershon RC, et al. (2013) NIH toolbox for assessment of neurological and behavioral function. Neurology 80(11 Supplement 3):S2–S6.

85. Gershon RC, et al. (2014) Language measures of the NIH toolbox cognition battery. J Int Neuropsychol Soc 20(6):642–651.

86. Fan J, McCandliss BD, Sommer T, Raz A, Posner MI (2002) Testing the efficiency and independence of attentional networks. J Cogn Neurosci 14(3):340–347.

87. Eriksen BA, Eriksen CW (1974) Effects of noise letters upon the identification of a target letter in a nonsearch task. Percept Psychophys 16.

88. Zelazo PD (2006) The Dimensional Change Card Sort (DCCS): A method of assessing executive function in children. Nat Protoc 1(1):297.

89. Salthouse TA, Babcock RL, Shaw RJ (1991) Effects of adult age on structural and operational capacities in working memory. Psychol Aging 6(1):118.

90. Bauer PJ, et al. (2013) III. NIH Toolbox Cognition Battery (CB): measuring episodic memory. Monogr Soc Res Child Dev 78(4):34–48.

91. Wechsler D (2014) Wechsler intelligence scale for children–Fifth edition (WISC-V): Technical and interpretive manual. Bloom MN Pearson Clin Assess.

92. Wulfert E, Block JA, Santa Ana E, Rodriguez ML, Colsman M (2002) Delay of gratification: impulsive choices and problem behaviors in early and late adolescence. J Pers 70(4):533–552.

93. Acker W, Acker C (1982) Bexley Maudsley automated processing screening and Bexley Maudsley category sorting test manual. Wind Engl NFER-Nelson.

94. Barch DM, et al. (2013) Function in the human connectome: task-fMRI and individual differences in behavior. Neuroimage 80:169–189.

95. Conley MI, et al. (2018) The racially diverse affective expression (RADIATE) face stimulus set. Psychiatry Res 270:1059–1067.

96. Tottenham N, et al. (2009) The NimStim set of facial expressions: judgments from untrained research participants. Psychiatry Res 168(3):242–249.

97. Logan GD (1994) On the ability to inhibit thought and action: A users’ guide to the stop signal paradigm. Psychol Rev 121(1):66–95.

98. Knutson B, Westdorp A, Kaiser E, Hommer D (2000) FMRI visualization of brain activity during a monetary incentive delay task. Neuroimage 12(1):20–27.

99. Yau W-YW, et al. (2012) Nucleus accumbens response to incentive stimuli anticipation in children of alcoholics: relationships with precursive behavioral risk and lifetime alcohol use. J Neurosci 32(7):2544–2551.

100. Ghasemi A, Zahediasl S (2012) Normality tests for statistical analysis: a guide for non-statisticians. Int J Endocrinol Metab 10(2):486–489.

101. Oba S, et al. (2003) A Bayesian missing value estimation method for gene expression profile data. Bioinformatics 19(16):2088–2096.

102. Dosenbach NUF, et al. (2017) Real-time motion analytics during brain MRI improve data quality and reduce costs. Neuroimage 161:80–93.

103. Hagler DJ, et al. (2018) Image processing and analysis methods for the Adolescent Brain Cognitive Development Study. bioRxiv:457739.

104. Fair DA, et al. (2018) Correction of respiratory artifacts in MRI head motion estimates. bioRxiv:337360.

105. Siegel JS, et al. (2014) Statistical improvements in functional magnetic resonance imaging analyses produced by censoring high-motion data points. Hum Brain Mapp 35(5):1981–1996.

106. Winkler AM, Webster MA, Vidaurre D, Nichols TE, Smith SM (2015) Multi-level block permutation. Neuroimage 123:253–268.

