## Supplementary Material for "Behavioral and neural signatures of working memory in childhood"

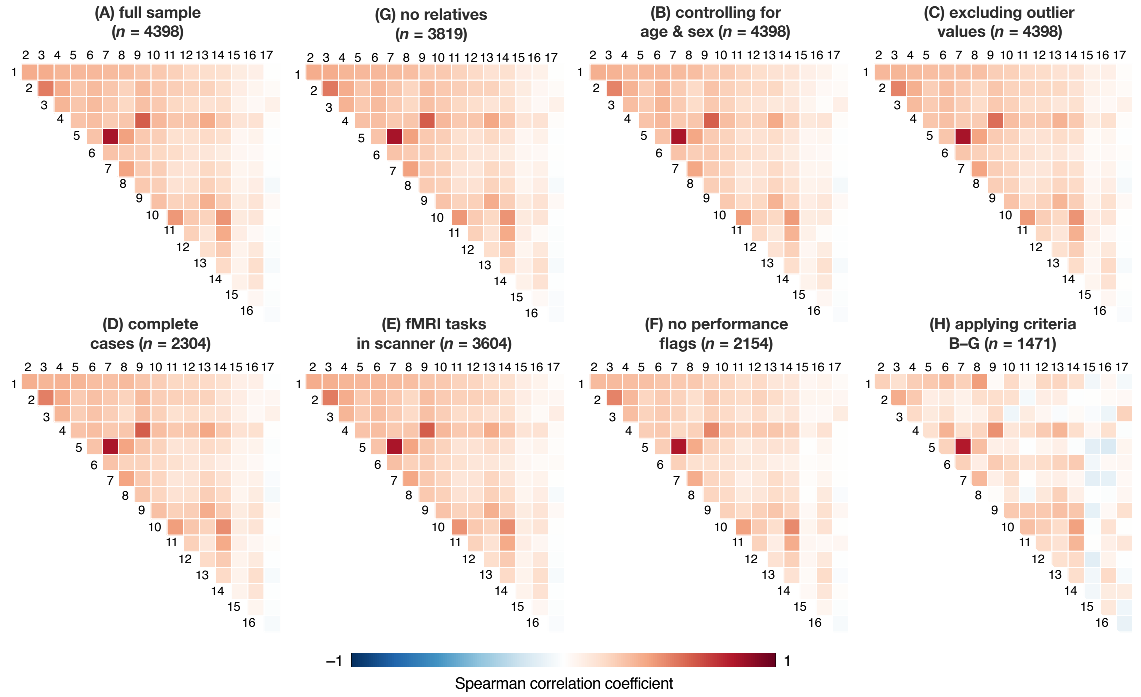


1. List Sorting Working Memory
2. Oral Reading Recognition
3. Picture Vocabulary
4. EN-back 2-back accuracy
5. RAVLT immediate recall
6. WISC-V matrix reasoning
7. RAVLT delayed recall
8. Picture Sequence Memory
9. EN-back 0-back accuracy
10. Dimensional Change Card Sort
11. Flanker
12. Little Man task efficiency ratio
13. EN-back recognition memory *d'*
14. Pattern Comparison
15. MID average earnings
16. SST –1*SSRT
17. Cash choice task

**Supplementary Figure 1**. Spearman correlations between performance measures in the full 4,398-child sample (A) and control samples (B–H). Measures are ordered according to the strength of their relationship with working memory, operationalized as NIH Toolbox List Sorting Working Memory Test, in the full sample.

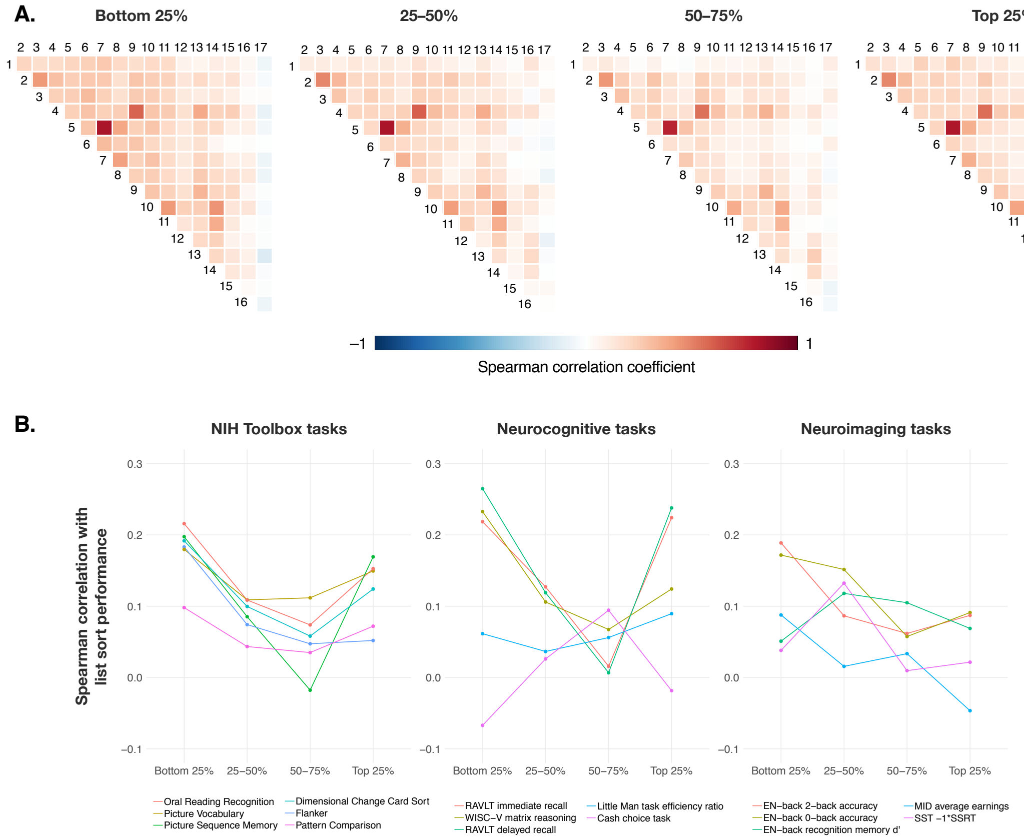


**Supplementary Figure 2.** (A) Spearman correlations between performance measures in children with the bottom 25% through the top 25% of list sorting scores. Each quartile includes data from 1086 children. (See Supplementary Figure 1 for a numbered list of behavioral measures.) (B) Relationships between working memory performance and other measures in each quartile of list sorting scores. The following comparisons reached statistical significance ($p <$ .05/96 comparisons): correlation between working memory and RAVLT delayed recall in the first vs. second quartile ($z = 3.53$, $p = 4.10\times10^{-4}$); correlations between working memory and picture sequence memory, RAVLT immediate and delayed recall, and matrix reasoning in the first vs. third quartile ($z$ values > 3.95; $p < 7.90\times10^{-5}$); correlations between working memory and picture sequence memory and RAVLT immediate and delayed recall in the third vs. fourth quartile ($|z|$ values > 4.39; $p < 1.13\times10^{-5}$); correlation between working memory and cash choice task selection in the first vs. third quartile ($|z| = 3.77$, $p = 1.66\times10^{-4}$).

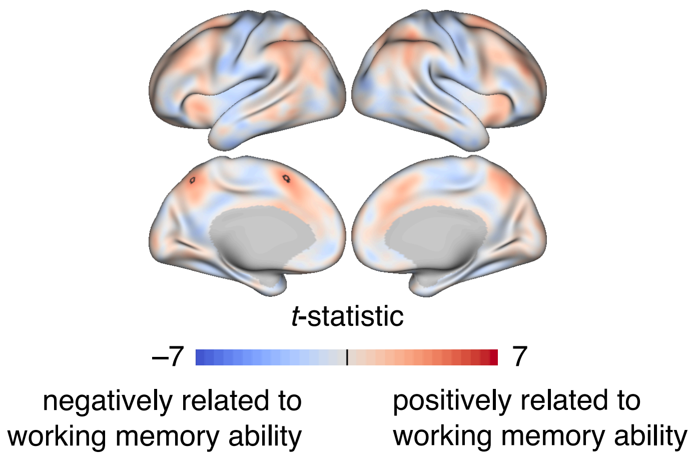


**Supplementary Figure 3.** Relationships between 2-back vs. 0-back activation and working memory function across individuals, controlling for age, sex, scanner, and in-scanner task accuracy (overall n-back accuracy, or *tfmri_nb_all_beh_ctotal_rate*; n = 1915). Black outlines indicate vertices significant at family-wise error-corrected, two-tailed p < .05. Results demonstrate that frontoparietal activation in 2-back vs. 0-back contrasts reflects trait-like in addition to state-like working memory abilities.

| | Description | Cognitive process(es) | Performance measure(s) | Data file | Variable name(s) | Percent missing | Percent outliers |
|---|---|---|---|---|---|---|---|
| Demo-graphics | Age in months, sex | | | abcddemo01 | *interview_age, gender* | 0 | |
| | Autism, epilepsy | | | abcd_screen01 | *scrn_asd, scrn_epls* | 0 | |
| | Family ID | | | acspsw02 | *rel_family_id* | 0 | |
| | Site ID | | | abcd_lt01 | *site_id_l* | 0 | |
| | Scanner ID | | | abcd_mri01 | *mri_info_deviceserialnumber* | 0.73 | |
| Neurocognitive battery | NIH Toolbox cognition battery | working memory | List Sorting Working Memory Test uss | abcd_tbss01 | *nihtbx_list_uncorrected* | 1.23 | 2.03 |
| | | language | Picture Vocabulary Test uss | abcd_tbss01 | *nihtbx_picvocab_uncorrected* | 0.93 | 1.65 |
| | | cognitive control, attention | Flanker Test uss | abcd_tbss01 | *nihtbx_flanker_uncorrected* | 1.00 | 2.18 |
| | | flexible thinking | Dimensional Change Card Sort Test uss | abcd_tbss01 | *nihtbx_cardsort_uncorrected* | 0.98 | 2.48 |
| | | processing speed | Pattern Comparison Processing Speed Test uss | abcd_tbss01 | *nihtbx_pattern_uncorrected* | 1.09 | .87 |
| | | visuospatial sequencing, memory | Picture Sequence Memory Test uss | abcd_tbss01 | *nihtbx_picture_uncorrected* | 1.02 | .74 |
| | | reading | Oral Reading Recognition Test uss | abcd_tbss01 | *nihtbx_reading_uncorrected* | 1.07 | 2.94 |
| | Matrix reasoning task | fluid reasoning | WISC-V matrix reasoning total scaled score | abcd_ps01 | *pea_wiscv_tss* | 5.37 | 1.18 |
| | Rey Auditory Verbal Learning Test (RAVLT) | learning, memory | total correct on immediate and delayed recall trials | abcd_ps01 | *pea_ravlt_sd_trial_vi_tc,*<br>*pea_ravlt_ld_trial_vii_tc* | 2.80<br>3.05 | 1.54<br>1.45 |
| | Intertemporal cash choice task | delay of gratification | choice of smaller-sooner or larger-later reward | cct01 | *cash_choice_task* | 1.55 | 0 |
| | Little Man task | mental rotation | efficiency ratio (% accuracy ÷ mean correct-trial RT) | lmtp201 | *lmt_scr_efficiency* | 4.82 | 1.34 |
| Functional MRI tasks | Emotional *n*-back (EN-back) task | memory, emotion regulation | % correct on 0-back and 2-back blocks | abcd_mri nback02 | *tfmri_nb_all_beh_c0b_rate,*<br>*tfmri_nb_all_beh_c2b_rate* | 28.33<br>28.33 | 3.30<br>3.11 |
| | Post-scan EN-back stimuli recognition memory test | memory | sensitivity (*d'*) | mribrec02 | mean of *tfmri_rec_all_beh_posf_dpr,*<br>*tfmri_rec_all_beh_neutf_dp,*<br>*tfmri_rec_all_beh_negf_dp,*<br>*tfmri_rec_all_beh_place_dp* | 31.63 | 1.43 |
| | Stop-signal task | impulsivity | −1*stop-signal RT | abcd_sst02 | *tfmri_sst_all_beh_total_meanrt* | 26.76 | 2.33 |
| | Monetary incentive delay task | reward processing | mean earnings | abcd_mid02 | mean of *tfmri_mid_r1_beh_t_earnings,*<br>*tfmri_mid_r2_beh_t_earnings* | 25.08 | 2.12 |

**Supplementary Table 1.** Demographic, neurocognitive, and neuroimaging task performance measures. Data were acquired from publicly available ABCD data release 1.1 (DOI 10.15154/1412097). Percent missing values represent the percentage of values missing in the full sample of 4,398 children meeting inclusion criteria, although note that recovery of missing data is ongoing. Percent outlier values represent the percentage of data values more than 2.5 standard deviations from the group mean. uss = uncorrected standard score.
